# Regulatory network structure and environmental signals constrain transcription factor innovation

**DOI:** 10.1101/2023.05.04.539253

**Authors:** Matthew J. Shepherd, Mitchell Reynolds, Aidan P. Pierce, Alan M. Rice, Tiffany B. Taylor

## Abstract

Evolutionary innovation of transcription factors frequently drives phenotypic diversification and adaptation to environmental change. Rewiring – that is gaining or losing connections to transcriptional target genes – is a key mechanism by which transcription factors evolve and innovate. However the frequency of functional adaptation varies between different regulators, even when they are closely related. To identify factors influencing propensity for rewiring, we utilise a *Pseudomonas fluorescens* SBW25 strain rendered incapable of flagellar mediated motility in soft-agar plates via deletion of the flagellar master regulator (*fleQ*). This bacterium can evolve to rescue flagellar motility via gene regulatory network rewiring of an alternative transcription factor to rescue activity of FleQ. Previously, we have identified two members (out of 22) of the RpoN-dependent enhancer binding protein (RpoN-EBP) family of transcription factors (NtrC and PFLU1132) that are capable of innovating in this way. These two transcription factors rewire repeatably and reliably in a strict hierarchy – with NtrC the only evolved rewiring route in a Δ*fleQ* background, and PFLU1132 the only evolved rewiring route in a Δ*fleQ*Δ*ntrC* background. However, why other members in the same transcription factor family have not been observed to rescue flagellar activity is unclear. Previous work shows that protein homology cannot fully explain this pattern, and mutations in rewired strains suggested high levels of transcription factor expression and activation drive rewiring. We predict that mutations that increase expression of the rewired transcription factor are vital to unlock rewiring potential. Here, we construct titratable expression mutant lines for 11 of the RpoN-EBPs in *P. fluorescens*. We show that in 5 additional RpoN-EBPs (HbcR, GcsR, DctD, AauR and PFLU2209), high expression levels result in different mutations conferring motility rescue, suggesting alternative rewiring pathways. Our results indicate that expression levels (and not protein homology) of RpoN-EBPs are a key constraining factor in determining rewiring potential. This suggests that transcription factors that can achieve high expression through few mutational changes, or transcription factors that are active in the selective environment, are more likely to innovate and contribute to adaptive gene regulatory network rewiring.

## Introduction

Propensity for innovation is a key determinant of evolvability – the ability to rapidly generate heritable phenotypic variation – in living organisms (Payne and Wagner, 2019). Alterations to transcription factor binding specificities and activity levels can facilitate rewiring of regulatory connections (Shepherd et al., 2022b; Taylor et al., 2022), which generate phenotypic variation and novelty that can provide selective advantages under changeable environmental conditions (Adhikari et al., 2021; Baumstark et al., 2015; Isalan et al., 2008; Martchenko et al., 2007; Patel and Matange, 2021). However propensity for engaging in rewiring varies between transcription factors (Igler et al., 2018), and whilst features influencing evolvability of transcription factor bindings sites have been investigated (Payne and Wagner, 2014; Tirosh et al., 2009), causes of variation in transcription factor evolvability remain poorly defined. To understand how novelty in regulatory systems evolves, we must identify intrinsic and environmental factors that determine rates of evolutionary innovation in these regulatory proteins.

A key property for evolvability in a transcription factor is its ability to rewire, which often involves gain of promiscuous activity. For a transcription factor, this constitutes gain of illicit or non-canonical regulatory interactions, and is a key factor in revealing a transcription factor to selection and for evolutionary innovation to occur (Alhindi et al., 2017; Copley, 2015). These interactions typically bare no physiological significance, but under the right selective conditions can become advantageous and drive innovation (Copley, 2020; Pougach et al., 2014). Previously, we investigated the evolutionary emergence of promiscuity and rewiring in the RpoN-dependant enhancer binding proteins (RpoN-EBPs) NtrC and PFLU1132 in the soil bacterium *Pseudomonas fluorescens* (Shepherd et al., 2022b; Taylor et al., 2015). We made use of an engineered maladapted GRN, where the RpoN-EBP flagellar master regulator FleQ is deleted resulting in loss of flagellar expression and motility. When challenged to rescue motility, NtrC and PFLU1132 evolved promiscuity and rewired to drive flagellar gene expression. Whilst this was in part due to the shared 3D structural homology between FleQ, NtrC and PFLU1132, which permits DNA binding without the need for mutation to the DNA binding domain, we also found that high gene expression and high levels of activation in these transcription factors was important for their promiscuous activity and rewiring (Shepherd et al., 2022b). These properties likely aid low-affinity promiscuous interactions to occur by providing an excess of a highly activated transcription factor that can saturate its native regulatory interactions and begin to engage in additional promiscuous regulatory activities.

The expression and activation levels of a transcription factor are determined by the architecture and connectivity of the GRN it sits within (Fang et al., 2017). Our previous findings therefore raise the possibility that pre-existing GRN architecture may significantly bias and constrain transcription factor evolution. For example, some transcription factors may face significant obstacles to gaining high expression levels. Many are negative autoregulators (41% of transcription factors in E. coli, Martínez Antonio et al., 2003), where multiple precise promoter mutations (Lamrabet et al., 2019) would be needed to increase expression level. Conversely, some transcription factors may be ideally suited to gaining high expression, through virtue of positive autoregulation (Bandyopadhyay and Banik, 2012; Goulian, 2010; Groisman, 2016), where loss of a negative repressor can easily lead to runaway feedback driving high transcription factor expression (diCenzo et al., 2017; Rao and Igoshin, 2021; Weyder et al., 2018).

Similarly, there will be variation in how easily a transcription factor may gain a hyperactivated state. This again depends on the signalling connectivity of the transcription factor within the GRN. Bacterial regulatory networks will sense and transduce internal or environmental stimuli (Seshasayee et al., 2006), commonly through two-component systems (TCS’s) where a sensor-kinase detects a signal and phosphorylates a REC receiver domain on a cognate transcription factor (Galperin, 2006). Alternatively, transcription factors can possess receiver domains that: directly bind small molecule signals, are bound by a protein inhibitor, or control sub-cellular localisation to determine activation (Browning et al., 2019). Mutations to TCS kinases are frequently observed to drive adaptation in regulatory systems (Mainiero et al., 2010; Moskowitz et al., 2012; Olaitan et al., 2014), and transcription factors that respond to particularly active or mutable cellular systems may more easily become hyperactivated.

In our model system, we have identified two RpoN-EBPs – NtrC and PFLU1132 – capable of rescuing flagellar motility via promiscuous activity. There are 19 other RpoN-EBPs that could also theoretically achieve this (Taylor et al., 2015) due to sharing structural homology with FleQ, however none are observed to do so in previous LB or M9 soft agar evolution experiments (Horton et al., 2021; Shepherd et al., 2022a, 2022b; Taylor et al., 2015a). We previously observed that a major constraining factor for PFLU1132 promiscuity was its gene expression level (Shepherd et al., 2022b), and as such we can hypothesise that gene expression level may be constraining evolutionary innovation through gain of promiscuity in the other RpoN-EBPs. To test this, we engineered titratable expression constructs for 11 of the RpoN-EBP genes encoded by *P. fluorescens* SBW25. This was achieved by introduction of the RpoN-EBP coding sequence downstream of a rhaSR-PrhaBAD L-rhamnose titratable promoter system (Meisner and Goldberg, 2016), and inserting this construct as a single copy into the *P. fluorescens* chromosome using the miniTn7 transposonal insertion system (Choi and Schweizer, 2006). This allows us to introduce a high level of expression for each RpoN-EBP into the GRN by addition of L-rhamnose to the growth media, simulating an alternative GRN architecture and environment where the transcription factor is highly expressed. Utilising these expression strains, we set out to identify if any other RpoN-EBPs could rescue motility, and if increased expression of an RpoN-EBP would bias evolutionary outcomes in a flagellar motility rescue experiment.

## Results

### Overexpression of RpoN-EBPs does not result in immediate rescue of motility in nearly all cases

We constructed rhamnose inducible expression systems for 11 *P. fluorescens* SBW25 RpoN-EBPs genes selected based on structural similarity to the FleQ protein, including representatives of orphan regulators (those lacking a cognate kinase) and members of two-component systems (Taylor et al., 2015b) – *aauR, algB, fleR, PFLU2055, PFLU1132, dctD* (*PFLU0286*), *prpR* (*PFLU2386*), *hbcR* (*PFLU2630*)*, gcsR* (*PFLU4895*), *mifR* (*PFLU4954*), *PFLU2695* and *PFLU2209.* These expression constructs were introduced into the *P. fluorescens* SBW25 Δ*fleQ*Δ*ntrBC* genetic background. This background lacks both the FleQ flagellar regulator, as well as the highly evolvable NtrBC system studied previously (Horton et al., 2021; Shepherd et al., 2022a; Taylor et al., 2015a) – preventing the presence of this dominant rewiring pathway from masking the effect of RpoN-EBP expression on evolutionary rescue of motility. Introduction of expression constructs was performed as single chromosomal insertions on a miniTn7 transposon. The native gene copies of the RpoN-EBPs were not deleted from the chromosome so this insertion results in effective duplication of the RpoN-EBP gene in questions. This allows conservation of the native regulatory network of each RpoN-EBP, with the expression construct simply acting to increase the concentration of the RpoN-EBP available within a cell without altering GRN structure.

We began by testing whether increasing the expression of each RpoN-EBP resulted in immediate restoration of motility. Constructs were incubated in M9 motility agar supplemented with 0.15% w/v L-rhamnose to induce expression. In almost all cases this did not result in immediate rescue of motility within 24 h. The exception was overexpression of *fleR* – an existing part of the flagellar regulatory cascade – which resulted in immediate flagellar motility (Supplementary Fig. 1A). This was curious, as there is no known mechanism for FleR to regulate the entire flagellar cascade by itself without the action of FleQ (Bouteiller et al., 2021; Dasgupta et al., 2003; Zhou et al., 2021), which may suggest a previously undiscovered regulatory connection (Supplementary Fig. 1B).

### Overexpression significantly increases the evolutionary rate of motility rescue for 5/11 RpoN-EBPs

To test if overexpression of each RpoN-EBP altered the mutational pathway for evolutionary rescue of motility, we incubated each expression construct in M9 motility agar with or without 0.15% Lrhamnose supplement for up to 6-weeks to allow motility mutants to evolve. In the Δ*fleQ*Δ*ntrBC* genetic background, the expected pathway of motility rescue is through the previously studied PFLU1131/2 two-component system (Shepherd et al., 2022). We measured the time taken for the evolution of a motile phenotype (termed “time to emergence”) in this background, as well as for each RpoN-EBP expression construct in the presence and absence of L-rhamnose. The Δ*fleQ*Δ*ntrBC* control background evolved with an average time to emergence that was not significantly different with or without L-rhamnose supplement (632 and 620 hours respectively, *P* = 0.8824 Wilcox test). Addition of L-rhamnose significantly reduced the time to emergence of motility across all RpoN-EBPs (Supplementary Fig. 1A). This was curious, as there is no known mechanism for FleR to regulate the entire flagellar cascade by itself without the action of FleQ (Bouteiller et al., 2021; Dasgupta et al., 2003; Zhou et al., 2021), which may suggest a previously undiscovered regulatory connection (Supplementary Fig. 1B).

### Overexpression significantly increases the evolutionary rate of motility rescue for 5/11 RpoN-EBPs

To test if overexpression of each RpoN-EBP altered the mutational pathway for evolutionary rescue of motility, we incubated each expression construct in M9 motility agar with or without 0.15% Lrhamnose supplement for up to 6-weeks to allow motility mutants to evolve. In the Δ*fleQ*Δ*ntrBC* genetic background, the expected pathway of motility rescue is through the previously studied PFLU1131/2 two-component system (Shepherd et al., 2022). We measured the time taken for the evolution of a motile phenotype (termed “time to emergence”) in this background, as well as for each RpoN-EBP expression construct in the presence and absence of L-rhamnose. The Δ*fleQ*Δ*ntrBC* control background evolved with an average time to emergence that was not significantly different with or without L-rhamnose supplement (632 and 620 hours respectively, *P* = 0.8824 Wilcox test). Addition of L-rhamnose significantly reduced the time to emergence of motility across all RpoN-EBPs (Fig.1), with mean time to emergence in the presence and absence of rhamnose being 356 and 674 hours respectively (Two-way ANOVA: F = 2.07e-12; P < 0.001). Broken down by each RpoN-EBP expression system, pairwise comparisons (Wilcox test) indicate no significant difference between presence and absence of L-rhamnose on time to emergence for *aauR, algB*, *PFLU2055*, PFLU2695, *mifR*, and *prpR*, but significant differences for *dctD*, *hbcR*, PFLU1132, PFLU2209, and *gcsR* (Fig. 1).

**Figure 1:**
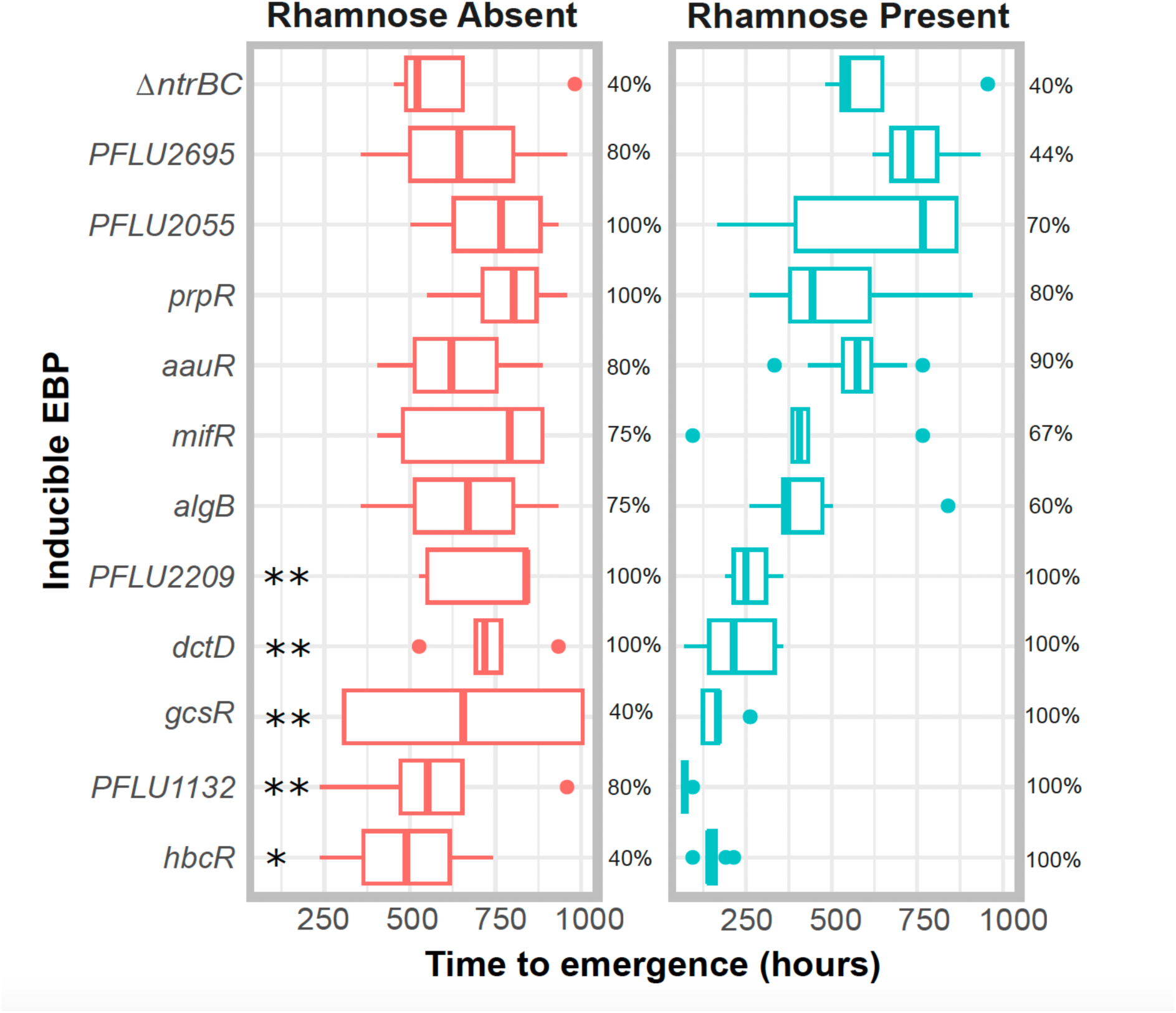
Time to emergence (TTE, hours) of flagellar motility in soft agar plates for each inducible RpoN-EBP system and the Δ*ntrBC* genetic background in the absence and presence of L-rhamnose (0.15% w/v). Boxplots display median, and upper and lower quartiles in standard format. Statistically significant differences in time to emergence are indicated by *’s with * = 0.005 < P < 0.05; ** = P < 0.005 (Wilcox test). To the right of each boxplot, a percentage is given indicating the proportion of independent replicate plates that evolved motility within 6 weeks.

### Overexpression of RpoN-EBPs results in switch of primary mutational targets, suggesting use of alternative evolutionary rewiring pathways

To identify whether increased RpoN-EBP expression had an effect on mutational targets utilised for evolutionary rescue of flagellar motility, we performed whole genome resequencing on isolates that evolved motility within the 6-week time frame for each evolution experiment. In the no rhamnose condition, the primary mutational target was the PFLU1130/1/2 operon, with mutations in the PFLU1130/1/2 locus being present in 100% of motile isolates for all RpoN-EBP expression strains tested in this media condition, as well as the *ΔfleQΔntrBC* control line (Fig. 2, Supplementary excel sheet E1). This was expected, as previous work identified this route as the primary pathway for evolutionary rescue of motility in the absence of *ntrBC* (Shepherd et al., 2022b). Many isolates had additional mutations alongside those in the PFLU1132 pathway, which are detailed in full in supplementary excel file E1. Two replicates in the no rhamnose control lines, one in an inducible *dctD* construct and one in an inducible *PFLU2209* construct, had mutations to *algB* alongside mutations in the PFLU1132 pathway – AlgB is an RpoN-EBP and FleQ homolog, so this may indicate involvement of this protein in rescuing motility in these isolates.

**Figure 2:**
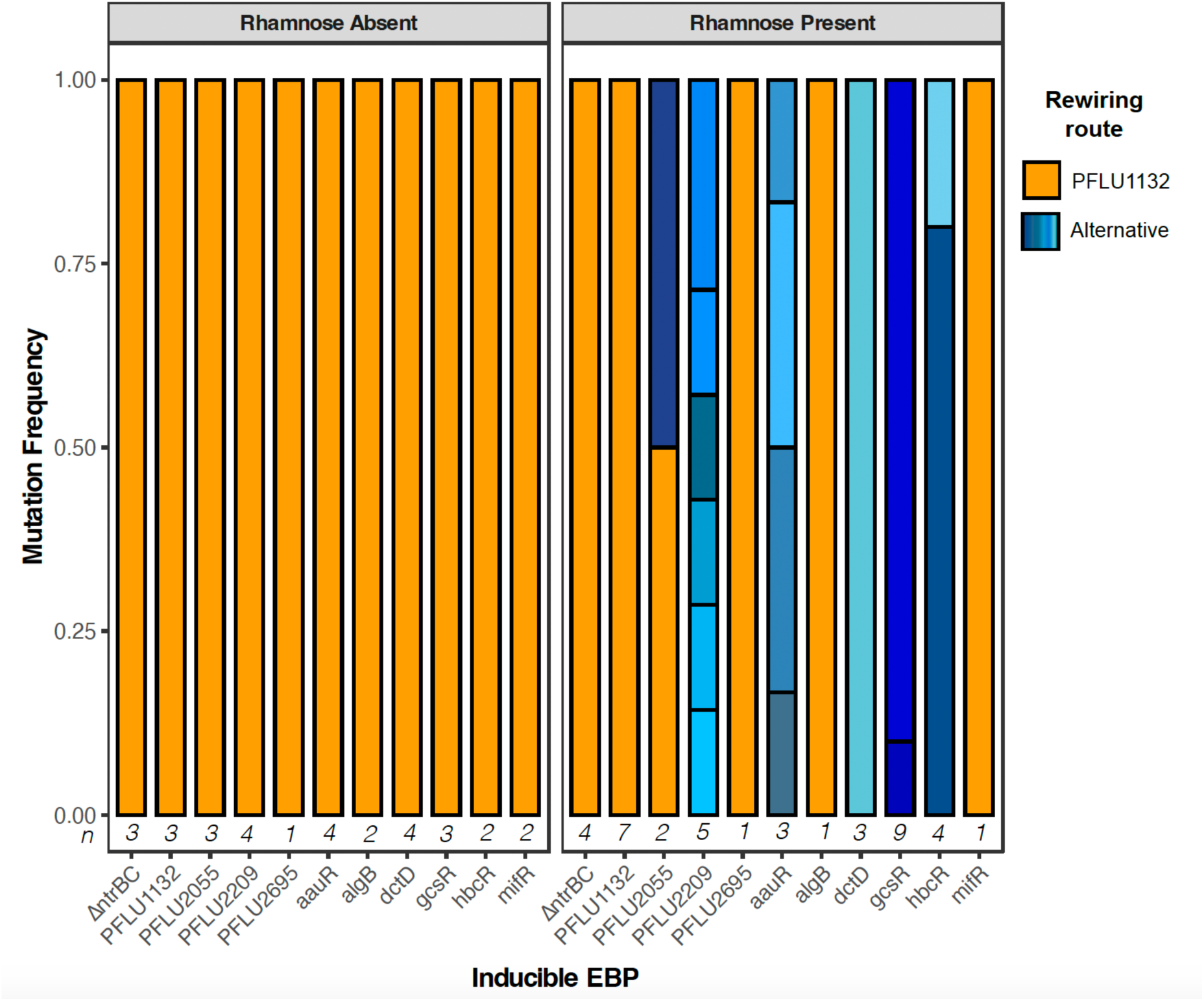
Proportion of observed mutations in evolved motile isolates. If a mutation was present in the PFLU1132 operon rewiring was assumed via this route, otherwise it is listed as an “alternative” rewiring route. Mutations in the absence (left) and presence (right) of L-rhamnose are shown in panels. The number of each independent RpoN-EBP expression lines that evolved motility and were subsequently sequenced in each experiment are given below each bar (*n*).

With addition of 0.15% w/v L-rhamnose to the soft agar, evolutionary rescue of motility via the PFLU1132 pathway became far less frequent, with the total number of isolates with PFLU1132 mutations across all RpoN-EBP expression conditions significantly dropping from 32/34 in the norhamnose control to 4/29 in the rhamnose condition (*x*^2^: P = 7.007e-10), with isolates from the *hbcR, aauR, gcsR, dctD* and *PFLU2209* overexpression conditions lacking any mutations in the PFLU1132 pathway. Additional control lines included a *ΔfleQΔntrBC* genetic background and an RpoN-EBP expression strain for PFLU1132 itself evolved in the presence and absence of L-rhamnose, as negative and positive controls respectively. For these controls, all motility rescue mutations in the presence of L-rhamnose occurred in the PFLU1132 pathway (Fig. 2 and 3).

**Figure 3.**
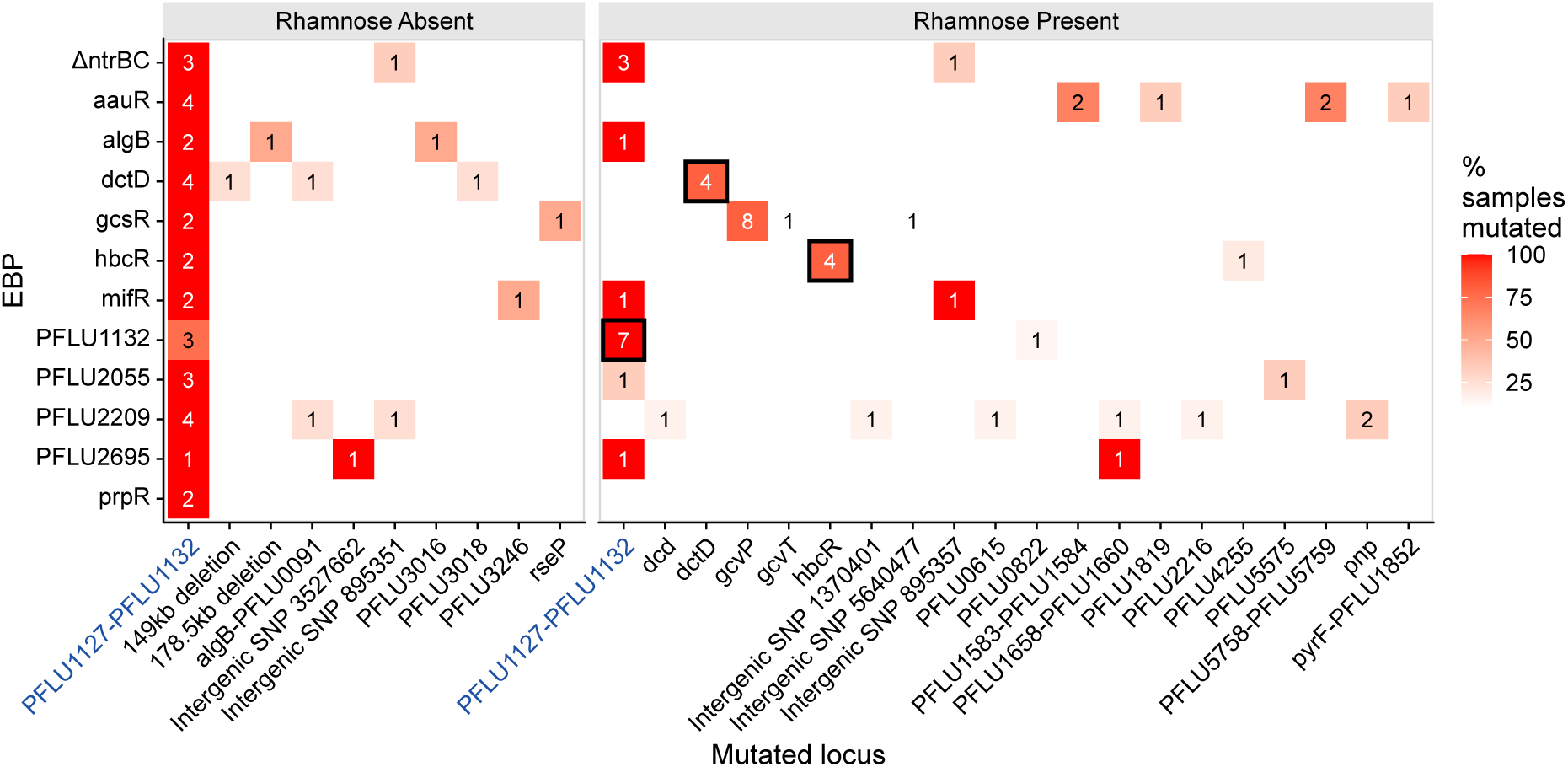
Counts of number of lines in which a genetic locus is mutated during motility rescue experiment for each RpoN-EBP expression line. Each panel is shaded by what percentage of the isolates in each condition have a mutation in that locus, e.g., where 100% of lines contain the same mutation, the box is coloured bright red. Note, some isolates have more than one mutation. Where the mutation is within a copy of overexpressed RpoN-EBP, the box has a black border. The primary ‘expected’ PFLU1132 pathway is indicated by a blue label on the x axis. The left panel shows the locus that was mutated in the motile isolates evolved in the absence of L-rhamnose, the right panel shows mutated loci in the motile isolates evolved in the presence of L-rhamnose.

In some cases, motile isolates evolved in the presence of L-rhamnose instead gained mutations associated with the relevant overexpressed RpoN-EBP. These could be grouped in to two broad categories. The first are those RpoN-EBP expression strains for which the primary mutation target switched to being one of the two copies of the RpoN-EBP being overexpressed. Overexpression of *hbcR*, *dctD* both resulted in rescue of motility predominantly with mutations to these transcription factor genes (Fig. 3). The second group do not gain mutations directly in the rhamnose-induced RpoN-EBP. These isolates instead gain mutations in a set of metabolic genes, which likely involve indirect modulation of RpoN-EBP activity through feedback.

The clearest example is that of *gcsR* overexpression, for which 8/10 motile isolates had gained a mutation in *gcvP*, 1 ins *gcvT* and 1 in the promoter of PFLU3504 (Fig. 3). Of the 8 *gcvP* mutations, 4 result in frame shifts, suggesting loss of function of the protein product. These genes encode the GcvP glycine dehydrogenase and the GcvT aminomethyltransferase, both components of the glycine cleavage system catabolic cycle (Okamura-Ikeda et al., 1993; Sarwar et al., 2016b). Loss of these catabolic enzymes will result in an accumulation of intracellular glycine, as seen previously for *gcvP* mutants in other organisms (Ernst and Downs, 2016). As GscR is a glycine responsive transcription factor (Sarwar et al., 2016), these mutations likely act to generate activating conditions for GcsR within the cytosol. Similarly, overexpression of *aauR* shifted the mutational spectrum to include genes involved in acidic amino acid metabolism. Across three motile isolates from the *aauR* condition there were individual cases of mutations in the genes *pyrB* (PFLU5758), *pyrC* (PFLU5759) and *pyrF* (PFLU1851) (Fig. 3). These genes encode enzymes aspartate carbamoyltransferase, dihydroorotase, and orotidine 5’-phosphate decarboxylase respectively, which are involved in the biosynthesis of pyrimidine nucleotides by converting aspartate to uridine-monophosphate (Schultheisz et al., 2011; West, 2010). Motile isolates from the *aauR* overexpression condition gained INDELs in these genes, which likely inactivated these enzymes (in the case of *pyrF*, a large portion of the open reading frame was deleted) – again this likely causes an increase of intracellular aspartate, by which AauR activity is determined through its sensor-kinase AauS (Yan et al., 2020). This would then generate conditions that activate the RpoN-EBP AauR in the cytosol. Interestingly, alongside mutations affecting aspartate catabolism, two motile *aauR* overexpression isolates had mutations in PFLU1583 and PFLU1584 (Fig. 3). These genes encode a putative anti-sigma factor system, which we have previously shown to facilitate RpoN-EBP promiscuity of PFLU1132 (Shepherd et al., 2022b). This suggests that mutations to PFLU1583/4 may constitute a general mechanism of enhancing promiscuity in RpoN-EBPs, as here mutations to this system are present in the absence of any PFLU1131/2 mutation. Finally, overexpression of *PFLU2055* and *PFLU2209* also resulted in motile isolates with putative feedback mutations. One *PFLU2055* motile isolate had a mutation to *lptD* (an LPS-assembly enzyme) – which may impact cell membrane integrity, which PFLU2055 homolog, PspF, has been shown to respond to (Joly et al., 2010). For PFLU2209, whilst its native function is unknown, mutations in motile isolates included genes involved in nucleotide metabolism (*pnp*, *dcd*) (Fig. 3).

Overexpression of these RpoN-EBPs resulted in significant shifts in mutational targets associated with rescue of motility, away from the primary PFLU1132 pathway utilised in these conditions. Rescue of motility primarily preceded through mutations acting on the overexpressed RpoN-EBP in question – either with mutation directly to the RpoN-EBP gene (as seen in *hbcR* and *dctD*), or the genes associated with its regulatory function, as was the case for *gcsR* and *aauR*. These results indicate that many members of the RpoN-EBP family of transcription factors are capable of rewiring to rescue motility in our assay, and that the expression of the RpoN-EBP is a significant factor constraining their potential to evolve novel interactions.

### Flagellar motility can be immediately rescued without mutation if an RpoN-EBP is overexpressed and activating environmental signals are present

We previously discussed the possibility that environmental conditions may ‘prime’ transcription factors for rewiring by providing conditions of high activation (Taylor et al., 2022).

Our experiments demonstrated that for some cases, an RpoN-EBP that was overexpressed could rescue motility through mutations that likely yielded increased cytosolic concentrations of the signal to which the RpoN-EBP in question responded (E.g., overexpression of glycine-responsive regulator *gcsR* resulted in mutations in the glycine cleavage pathway). Such activating signals could also feasibly be provided by the external environment rather than mutation to internal metabolism. To test this, we selected two RpoN-EBPs for which an activating signal could be provided externally – *gcsR*, and *hbcR*. This pair were chosen as contrasting examples – *gcsR* rescued motility in our experiment through mutations to internal glycine catabolism, whereas *hbcR* evolved through mutation to one of the two copies of the RpoN-EBP gene itself. However, the HbcR transcription factor can be activated by the presence of (R)-3-hydroxybutyrate (R-HB), a metabolite which cannot be produced by the internal metabolism (Lundgren et al., 2015). This may explain why in our assay, mutation to *hbcR* itself are the primary mechanism of rescuing motility for this RpoN-EBP, as higher levels of R-HB cannot be produced internally.

To test if flagellar motility could be rescued by environmentally providing activating signals for these two transcription factors, we set up M9 soft agar plates supplemented with L-glycine for *gcsR* or RHB for *hbcR*. In both cases, incubation of these strains in the presence of both L-rhamnose (to ensure RpoN-EBP overexpression), and the relevant activating signal resulted in immediate flagellar motility (Fig 4). We additionally demonstrated that activating signals that facilitate rewiring can be provided environmentally by overexpressing the *ntrBC* two component system in the presence of glutamate, which resulted in immediate motility – curiously this did not result in motility when only *ntrC* was overexpressed – which likely indicates the importance of TCS stoichiometry (Supplementary Fig. 1A). This was curious, as there is no known mechanism for FleR to regulate the entire flagellar cascade by itself without the action of FleQ (Bouteiller et al., 2021; Dasgupta et al., 2003; Zhou et al., 2021), which may suggest a previously undiscovered regulatory connection (Supplementary Fig. 1B).

**Figure 4:**
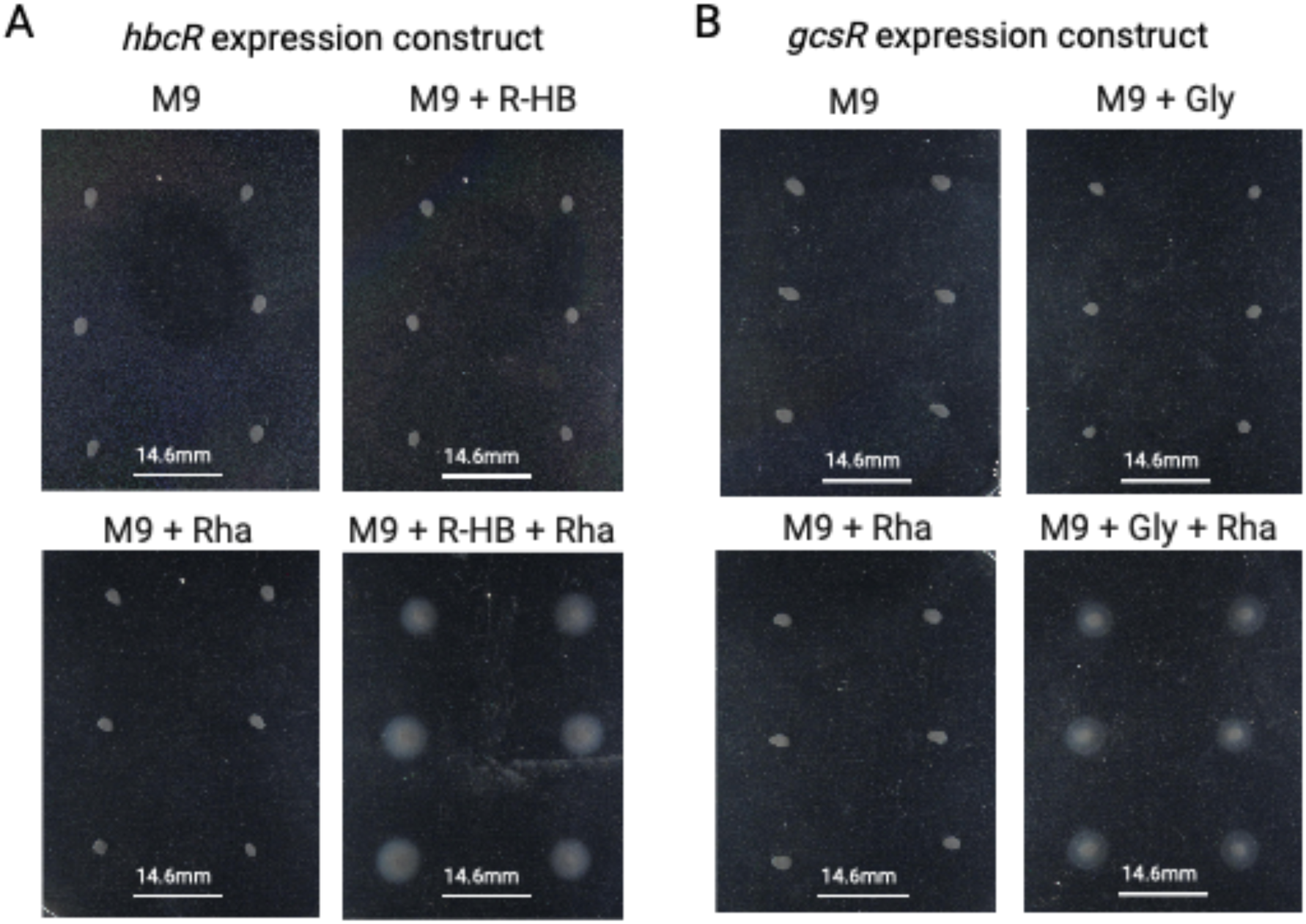
Flagellar motility can be rescued by alternative RpoN-EBPs if overexpressed and in the presence of an activating signal. **A)** *hbcR* and **B)** *gcsR* after 24hrs of incubation in 0.25% agar motility plates. Plates consisted of M9 media, with either no supplement, 0.15% L-rhamnose (+Rha), an environmental signal for each RpoN-EBP (30mM R-hydroxybutyrate (+R-HB) for *hbcR*, or 10mM Glycine (+Gly) for *gcsR*), or both signal and L-rhamnose.

To confirm that motility depends on overexpression of these RpoN-EBPs and their interaction with their respective signals, control plates of the rhamnose-inducible expression system lacking any RpoN-EBP, and of the *mifR* overexpression strain (RpoN-EBP overexpressed should not respond to either signal) were run (Supplementary Fig S3) and were found to remain immotile upon addition of L-rhamnose, R-HB and L-glycine in the relevant combinations. This confirms that rescue of flagellar motility via rewiring of RpoN-EBP transcription factors can also be facilitated by presence of signals that activate the relevant transcription factor in the environment.

## Discussion

Through manipulation of transcription factor expression, we have identified several members of the RpoN-EBP protein family capable of rewiring to rescue flagellar mediated motility in *P. fluorescens*, in addition to those that have been previously identified to rewire without modified expression (NtrC and PFLU1132) (Shepherd et al., 2022b; Taylor et al., 2015a). Regulators HbcR, AauR, GcsR, DctD, and PFLU2209 all became viable rewiring candidates upon induced high expression, either via mutation to the transcription factor itself (HbcR, DctD), or via physiological feedback (AauR, GcsR, PFLU2209). These results lend significant strength to our prior hypothesis that the expression of a transcription factor (and by extension its connectivity within the gene regulatory network) will constrain its ability to gain promiscuous activity and undergo rewiring (Shepherd et al., 2022b). Additionally, we have previously suggested that hyperactivation of a transcription factor is important for rewiring (Shepherd et al., 2022b), and again our results support this – HbcR, GcsR and AauR have all been shown to be capable of rescuing motility through production (or external supply) of an activating metabolic signal.

This set of identified rewirable transcription factors, together with NtrC and PFLU1132, constitute 58% of the RpoN-EBP transcription factor family tested by our work, and 33% of the total encoded by *Pseudomonas fluorescens* SBW25. For all these regulators, no canonical regulatory link to the flagellum is known, yet they are capable of driving flagellar expression. None of these RpoN-EBPs are the most structurally homologous to FleQ (Taylor et al., 2015b), so it is quite clear that whilst structural homology to FleQ matters (no non-homologs have been rewired in evolution assays), protein homology to the typical regulator of a gene is not a key factor in deciding whether a particular regulator undergoes innovation to rewire and gain a novel connection to that gene. A caveat to this result is that we currently only consider structural homology across the entire protein. It could be that homology within the DNA-binding domain is a better predictor of promiscuous binding potential and this is an active and exciting area of future research.

The fact that so many members of the RpoN-EBP transcription factor complement in *P. fluorescens* SBW25 can rescue flagellar motility suggests that this transcription factor family may be uniquely suited to regulatory innovation. Why this should be the case is unclear, however the answer may lie in the unique regulatory mechanism of the RpoN-EBP family. The sigma factor, RpoN, plays a pivotal role in localising the EBP so that it may provide ATPase activity to catalyse transcription initiation (Doucleff et al., 2007). RpoN-EBPs are indeed a highly diverse and varied class of transcription factors (Bush and Dixon, 2012), and numbers of RpoN-EBPs encoded can significantly vary between bacteria: *P. fluorescens* encodes ∼20*, Escherichia coli* ∼12, *Bacillus subtilis* ∼5, *Chlamydia pneumoniae* and *Treponema pallidum* both encode 1 each, and *Mycoplasma genitalium* and *Rickettsia prowazekii* encode none (Studholme and Buck, 2000). Even within a single order or genus, the number can be highly variable, with a range of 1-35 RpoN-EBPs present in 57 *Clostridiales* species (Nie et al., 2019), and between 9-29 RpoN-EBPs within the *Pseudomonas* genus (see methods, Supplementary excel sheet E2).

It has been suggested previously that RpoN-EBPs provide a biological advantage in regulating genes compared to other regulators, potentially due to providing ‘leaky’ regulation (Studholme and Buck, 2000). This can be an advantage in changeable environments (Schmutzer and Wagner, 2020), situations also evidenced to drive regulatory rewiring and innovation within GRNs (Tsuda and Kawata, 2010). Our results indicate that this family of regulators may be well suited toward regulatory innovation, a key factor in coping with changeable conditions. Additionally, our findings suggest that possessing large sets of transcription factors derived from the same protein family increases the opportunity for promiscuity and innovation to occur via alteration to interconnectivity between similar regulatory proteins.

Alongside showcasing the ability of a set of RpoN-EBPs to rewire when overexpressed, we have also demonstrated that signals that activate transcription factors can also aid rewiring. The nature of connections within regulatory networks will not only determine which transcription factors are highly expressed but also which are activated at any given moment via transduction of environmental signals (Friedlander et al., 2016; Jothi et al., 2009). Both the pre-existing GRN architecture and the prevailing environmental conditions both therefore have the potential to influence which transcriptional regulators are rewirable and evolvable. We have previously suggested that environmental conditions may ‘prime’ particular transcription factors for rewiring (Taylor et al., 2022). Our results in this work support this suggestion, as we demonstrate that activating signals can result in rescued flagellar motility when present in combination with increased transcription factor expression for regulators HbcR and GcsR.

Our data also raise the intriguing possibility that transcription factors that are responsive to signals, which can be generated via mutation to internal cellular physiology, may more easily evolve to rewire than those than respond to purely external signals. In the case of AauR and GcsR, mutations occurred in metabolic pathways that catabolise the chemical signals to which these regulators respond (Aspartate and Glycine), likely resulting in their activation. Similarly, in our previous studies of NtrBC rewiring, the *glnA* gene encoding glutamine synthetase was a possible mutational target (Horton et al., 2021; Shepherd et al., 2022; Taylor et al., 2015). Loss of function to GlnA generates a condition of low glutamine concentration within the cell, which triggers activation of NtrBC through GlnK (Hervás et al., 2009). This type of internal metabolic mutation that will yield a high concentration of an activating metabolite signal did not occur for HbcR – mutations primarily targeted the transcription factor itself – possibly because its activating signal, R-HB, cannot be generated by internal metabolism (Lundgren et al., 2015). When R-HB was supplied externally alongside high HbcR expression, motility was restored. The ease by which such an activating signal is provided is therefore also important in shaping and constraining the ability of a transcription factor to rewire, alongside the expression level of that transcription factor. Regulators that respond to external physical conditions (e.g., light or temperature), or exogenous metabolites (e.g., R-HB) may therefore be less likely to engage in evolutionary rewiring.

Our results indicate the key role that GRN architecture plays in constraining avenues of evolutionary innovation and evolvability in their constituent transcription factors. We have demonstrated that by increasing the expression of a transcription factor – representing an altered GRN architecture – that transcription factor can become the preferred candidate for evolutionary rewiring in our model system. We have demonstrated propensity for rewiring in multiple RpoN-EBP family members and highlighted the interplay between GRN architecture and prevailing environmental conditions necessary to facilitate rewiring in certain transcription factors. Our findings help to build a systems level understanding of how conditions and signalling within gene regulatory networks can influence the evolutionary trajectories of their constituent components and create or constrain opportunities for innovation of transcription factors.

## Acknowledgements

This work was funded by a Royal Society Research Fellows Grant (RG160491; awarded to TBT) supporting MJS; Windsor Fellowship Syncona PhD Scholarship (awarded to MR) supporting MR; Royal Society Research Fellows Enhancement Award (RF\ERE\210249; awarded to TBT) supporting AMP; Royal Society Dorothy Hodgkin Research Fellowship (DH150169; awarded to and supporting TBT).

---

**Supplementary Figure S1:**
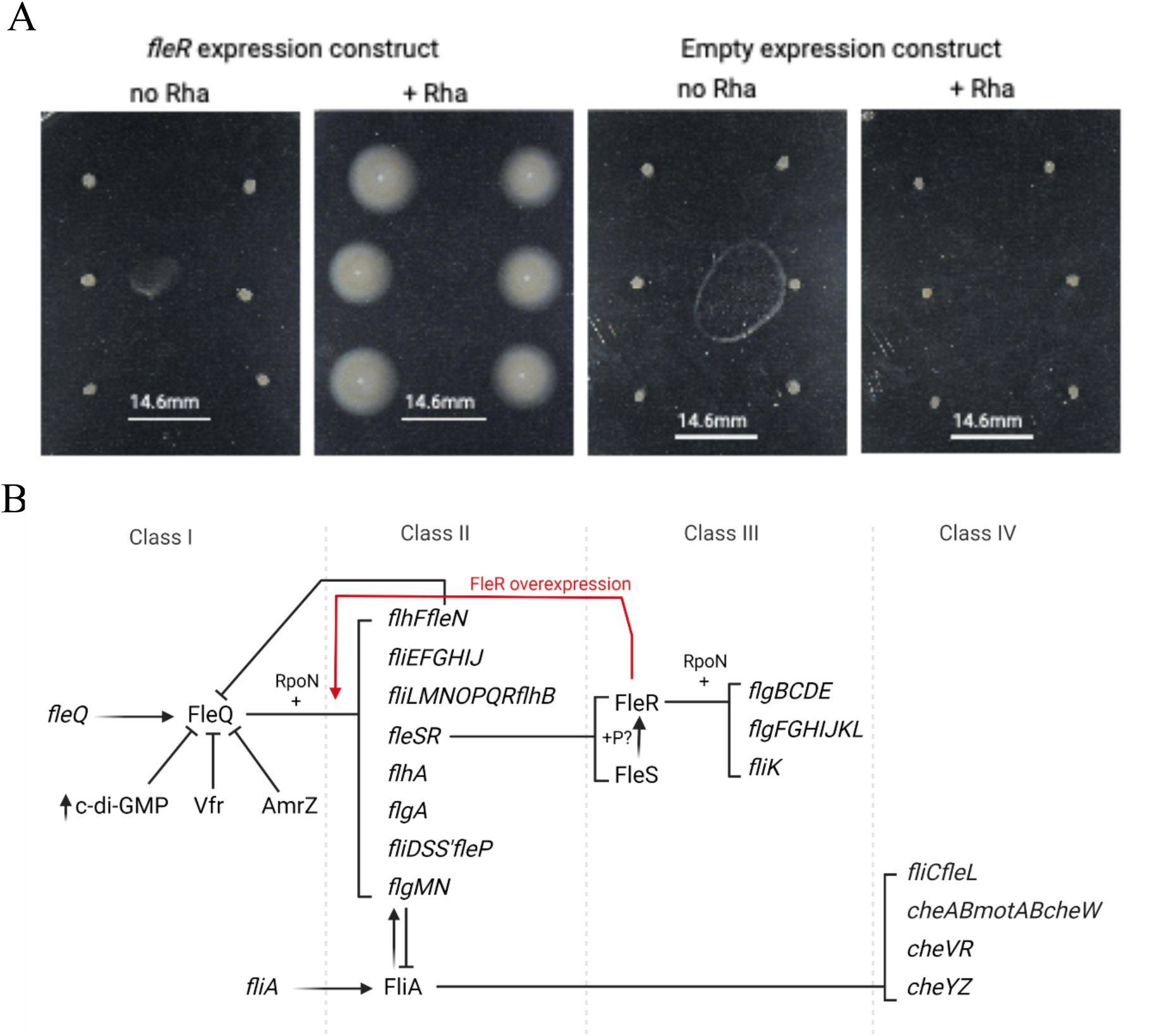
Overexpression of RpoN-EBP FleR grants flagellar motility. **A)** Images of *fleR* overexpression strain in LB plates with and without L-rhamnose supplement (0.15% w/v), and vector-only control lacking an RpoN-EBP coding sequence. **B)** Diagram of flagellar hierarchical regulatory cascade. Putative FleR regulatory connection identified in this experiment is shown in red. Adapted from published works (Bouteiller et al., 2021; Dasgupta et al., 2003).

**Supplementary Figure S2:**
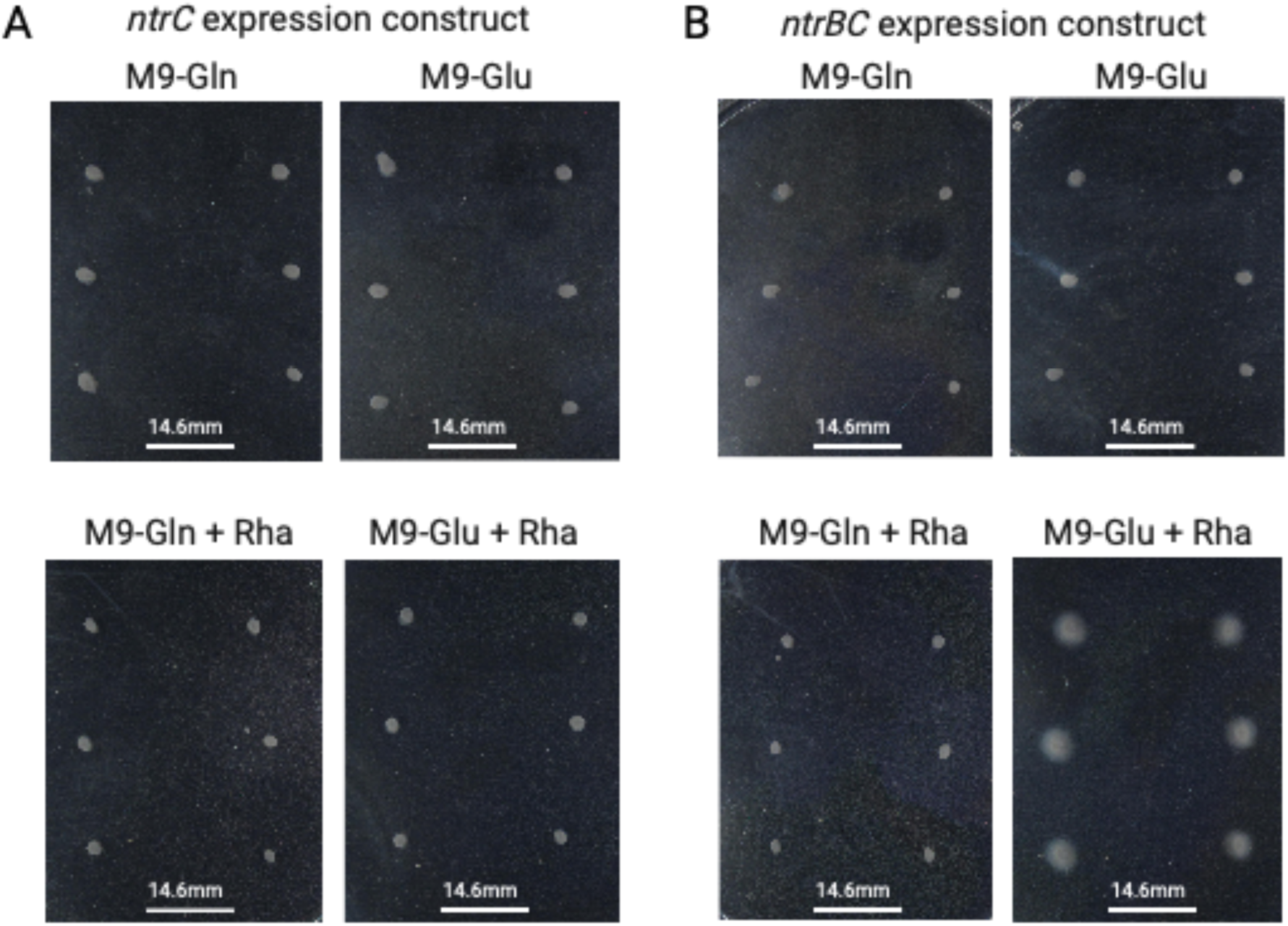
Overexpression of *ntrBC* but not *ntrC* alone in presence of activating conditions results in immediate flagellar motility. AR2 miniTn7[rha-ntrC] and AR2 miniTn7[rha-ntrBC] after 24hrs of incubation in 0.25% agar motility plates. Plates consisted of M9 media with either glutamine (Gln – does not activate *ntrBC*) or glutamate (Glu – activates *ntrBC*) as sole nitrogen sources, with or without 0.15% L-rhamnose (+Rha)

## Materials and Methods

### Strains and culture conditions

All ancestral strains in this study are derived from either *Pseudomonas fluorescens* AR2 (SBW25Δ*fleQ* IS-ΩKm-hah: PFLU2552), or AR2ΔntrBC (constructed previously, Shepherd et al., 2022). AR2 lacks flagellar master regulator FleQ and possesses a transposon-insertional disruption of the gene *viscB* (PFLU2552), rendering it immotile as detailed previously (Alsohim et al., 2014; Taylor et al., 2015a). All routine culturing of strains were cultured on lysogeny broth (LB; Miller) media at 27°C. *Escherichia coli* strains for cloning were cultured on LB media at 37°C. For experiments using M9 minimal media, the follow recipe was used: 0.2% w/v Glucose, 0.1mM CaCl_2_, 2mM MgSO_4_, 1x M9 salts (33.7 mM Na_2_HPO_4_, 22mM KH_2_PO_4_, 8.55 mM NaCl, 9.35 mM NH_4_Cl). In some cases, alternatives were used as the sole carbon or nitrogen sources, in which the Glucose or NH_4_Cl were omitted respectively.

### RpoN-EBP expression system construction

Inducible expression constructs for a panel of RpoN-EBPs were constructed in the AR2 genetic background. The RpoN-EBP ORF was amplified by PCR and a strong ribosome binding site (stRBS) introduced upstream with a 7bp short spacer between the stRBS and the start codon. PCR also introduced restriction enzyme cut sites up and down-stream of the gene, which were used to insert the RpoN-EBP into the multiple cloning site of the miniTn7 suicide vector pJM220 (obtained from the Addgene plasmid repository, plasmid #110559) by restriction-ligation. This positions the RpoN-EBP gene under control of the PrhaBAD promoter and downstream of rhaSR, allowing rhamnose-titratable expression of the RpoN-EBP (Meisner and Goldberg, 2016) as used to do so previously (Shepherd et al., 2022). This construct was transformed into *E. coli* DH5α by chemical-competence heat-shock. The miniTn7 transposon containing the *rhaSR* genes and the P*rhaBAD*-stRBS-RpoN-EBP construct was then transferred to the *P. fluorescens* chromosome by transposonal insertion downstream of the *glmS* gene via four-parent puddle-mating conjugation (Choi and Schweizer, 2006). The relevant *E. coli* DH5α pJM220-derived plasmid donor was combined with recipient *P. fluorescens* AR2 strains, transposition helper *E. coli* SM10 λpir pTNS2 and conjugation helper *E. coli* SP50 pRK2073, and Gentamicin resistant *Pseudomonas* selected for on LB supplemented with Gentamicin sulphate and Kanamycin sulphate. Chromosomal insertion of the correct miniTn7 transposon and RpoN-EBP was confirmed by colony PCR. The *rhaSR*-P*rhaBAD*-stRBS-PFLU4895 construct was also transferred to the chromosome by miniTn7 insertion in an AR2Δ*ntrBC* background. Activity of the rhaSR-PrhaBAD and response to L-rhamnose present in growth media tested using a *rhaSR*-P*rhaBAD*- stRBS-*lacZ* construct in a β-galactosidase activity assay as presented previously (Shepherd et al., 2022b).

### Motility rescue evolution experiments

Evolutionary rescue of motility was assayed in 0.25% agar M9 plates as described previously (Horton et al., 2021; Taylor et al., 2015a). Pure single colonies were picked and inoculated using a sterile toothpick and incubated at 27°C. Plates were checked a minimum of twice daily for motility, recording time to emergence. Motile zones were sampled immediately and always from the leading edge. Motile isolates were streaked on LB agar, and a pure colony picked and stored at −80°C as glycerol stocks of LB overnight cultures. All subsequent analysis was conducted on these pure motile isolates. Experiment was run for six weeks and any replicates without motility after this cut-off recorded as having not evolved.

### Mutation identification by whole genome resequencing

To identify motility rescuing mutations, genomic DNA was extracted from motile strains and their ancestral strain using the Thermo Scientific GeneJET Genomic DNA Purification Kit. Genomic DNA was quality checked using BR dsDNA Qubit spectrophotometry to determine concentration and nanodrop spectrophotometry to determine purity. Illumina NextSeq 2000 sequencing was provided by Seqcenter (Pittsburgh, PA, USA), with a minimum 30x coverage. Returned paired-end reads were further filtered using fastp v0.23.2 (Chen et al., 2018) with parameters --disable_adapter_trimming -- cut_front --cut_tail --cut_mean_quality 30 --qualified_quality_phred 30 --length_required 50.

Alignment of quality trimmed reads to the *P. fluorescens* SBW25 reference genome (Silby et al., 2009) and mutation identification was conducted using breseq (Deatherage and Barrick, 2014). Four substitutions/indels were observed in 92-100% of samples and were ignored in downstream analysis as these mutations were assumed to be present in the AR2 genetic background with respect to the SBW25 reference genome. These mutations were: three small indels at positions 45881 (86/93 samples), 985333 (93/93), 3447984 (92/93), and an intergenic substitution at position 1786536 (93/93). Five SNPs present in both ancestral strains and evolved motile strains are listed in Supplementary excel sheet E1 but not included in Figure 3. Additionally, two samples gcsR_C1 and ΔntrBC_1 have a mutS mutation and subsequently far more additional mutations relative to other samples. These mutations are listed in Supplementary excel sheet E1 but not included in Figure 3.

### RpoN-EBP protein domains

To identify where mutations affect the protein domains of RpoN-EBPs, the *P. fluorescens* SBW25 proteome was searched for Pfam 35.0 domains (Mistry et al., 2021) using InterProScan v5.59-91 (Jones et al., 2014).

To count RpoN-EBPs in *Pseudomonas* species, complete *Pseudomonas* genomes were obtained from NCBI RefSeq (O’Leary et al., 2016). A representative strain was selected for species with numerous strains. After ensuring ≥90% of *Pseudomonadales* marker genes were present and ≤2% were duplicated using BUSCO (Manni et al., 2021), 140 genomes remained. The proteomes were searched using InterProScan for “Sigma-54 interaction domain (PF00158)” Pfam signature to identify RpoN-EBPs.

### Induced motility experiments

Motility phenotype induction was assayed using soft-agar (0.25%) plates with varying media compositions and supplements. Soft-agar plates were set up as described previously (Alsohim et al., 2014). For induction of the *rhaSR*-P*rhaBAD* RpoN-EBP expression constructs, L-rhamnose was added to the motility agar with a final concentration of 0.15% w/v. Activating signals R-hydroxybutyrate (R-HB-for *hbcR*), or Glycine (Gly-for *gcsR*) were added to media at concentrations of 30mM and 10mM respectively. Six biological replicates of each strain of interest were inoculated into the motility plates by picking single colonies with a sterile toothpick and stabbing into the soft agar. These replicates allow us to discern whether any motility is adaptive, as if all 6 replicates display the same phenotype it is unlikely that any mutations have occurred.

### Statistical analysis

All statistical analysis and data handling was performed using R core statistical packages. Differences between the rhamnose absent and present conditions were tested via two-way ANOVA, and differences between the two conditions for individual RpoN-EBPs tested using Wilcoxon tests. For differences in rewiring pathway frequency between rhamnose absent and present conditions, a *x*^2^ test was used.

**Supplementary Table T1:**
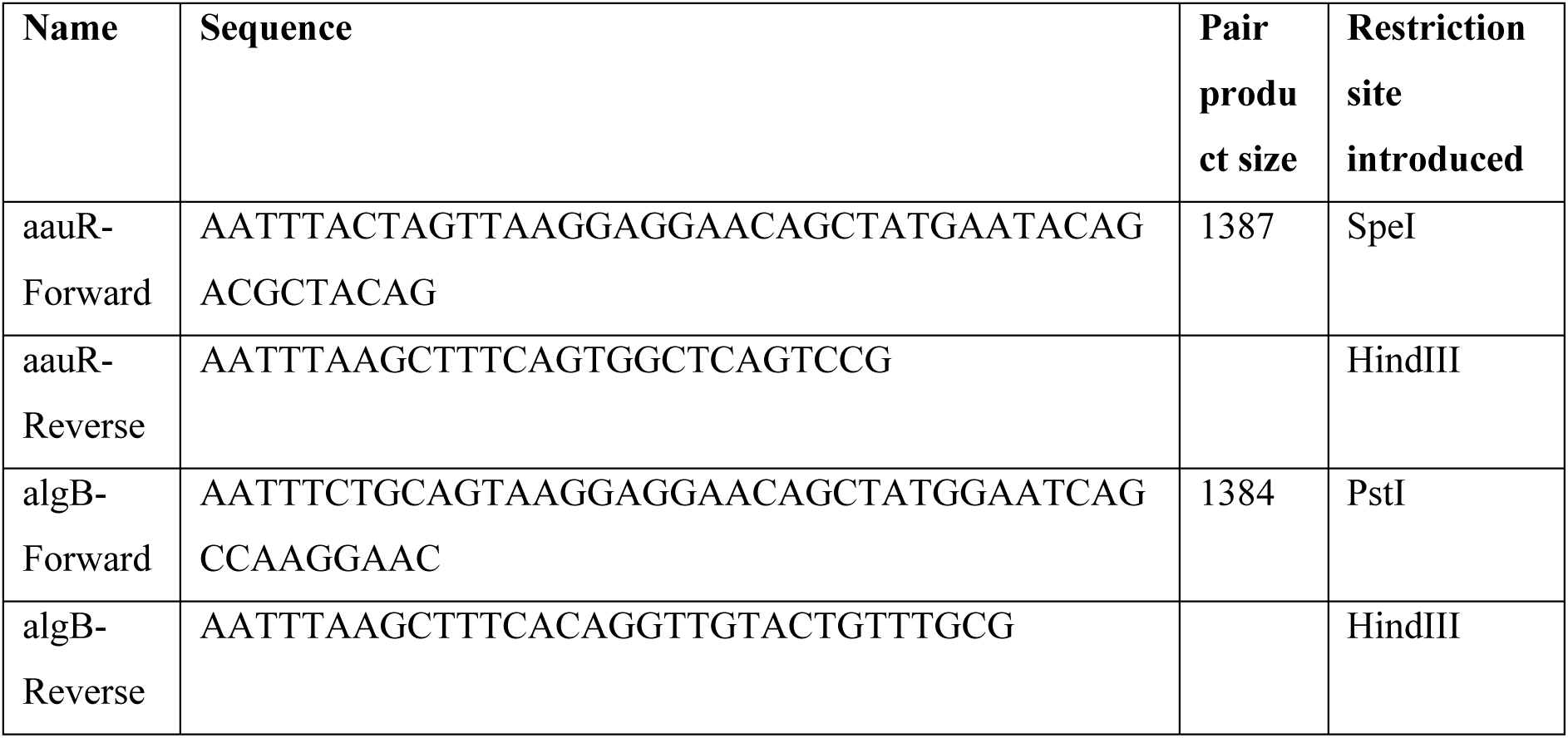
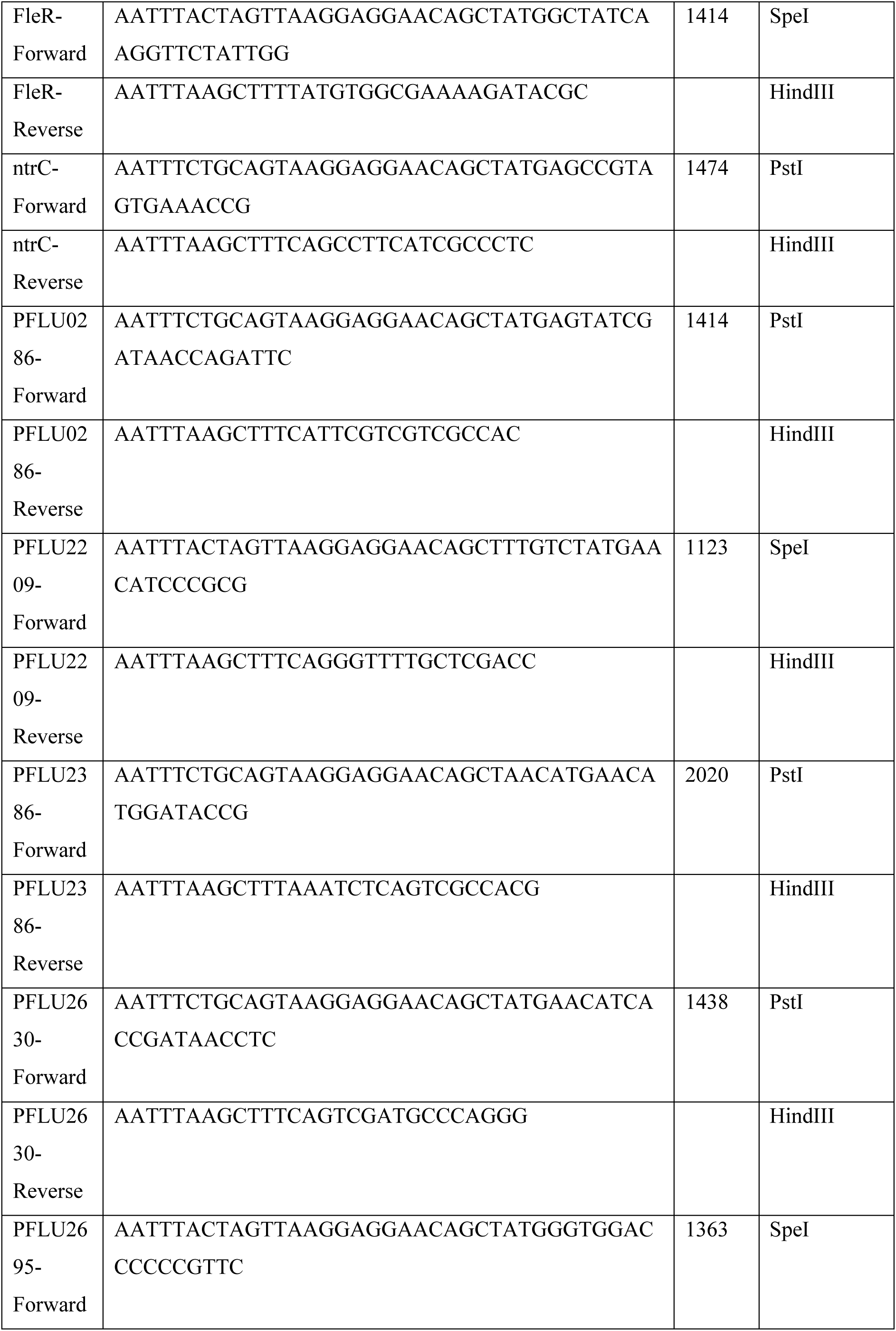
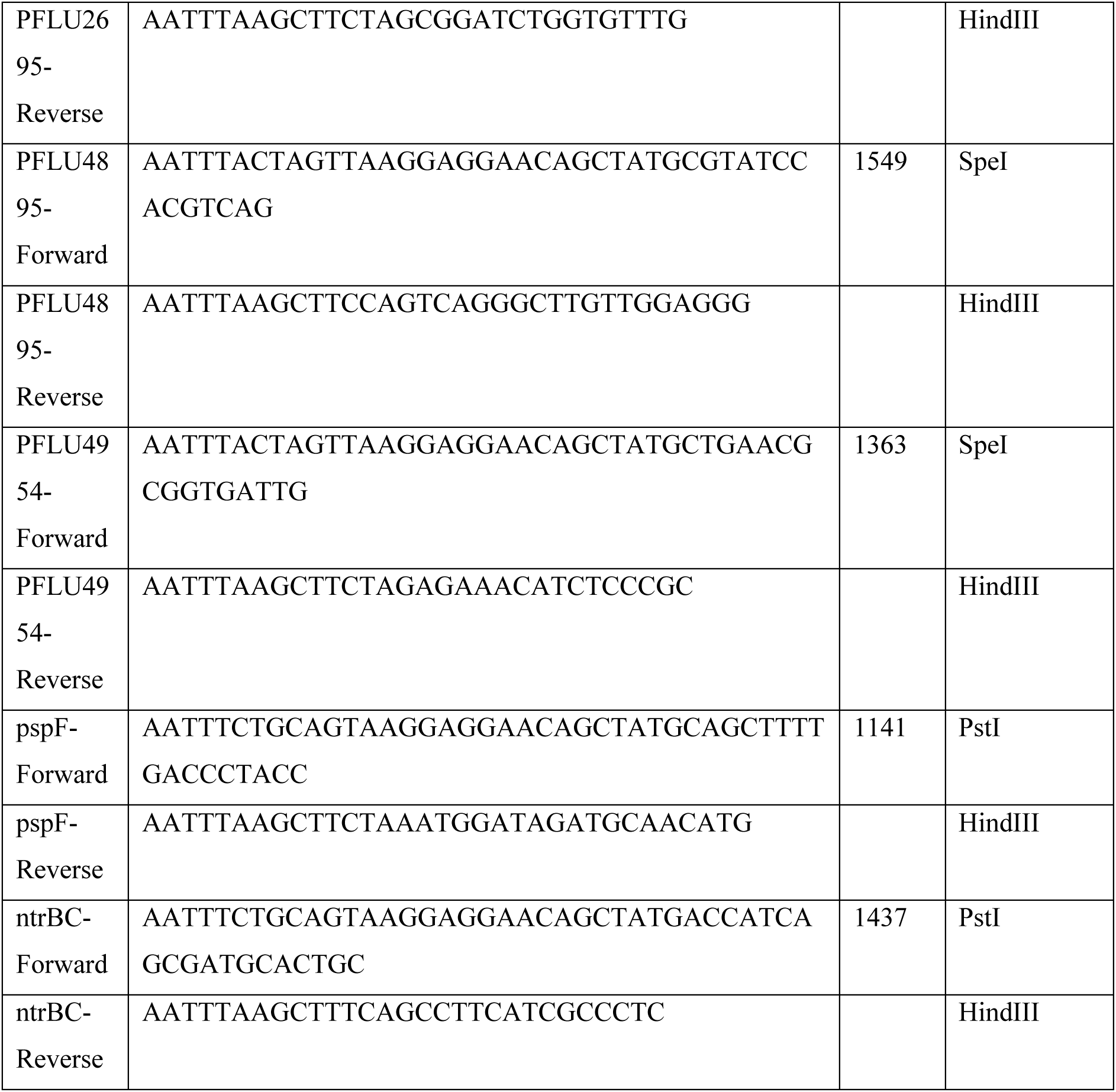
Forward and reverse primers used for amplification and construction of mutants.

**Supplementary Figure S3:**
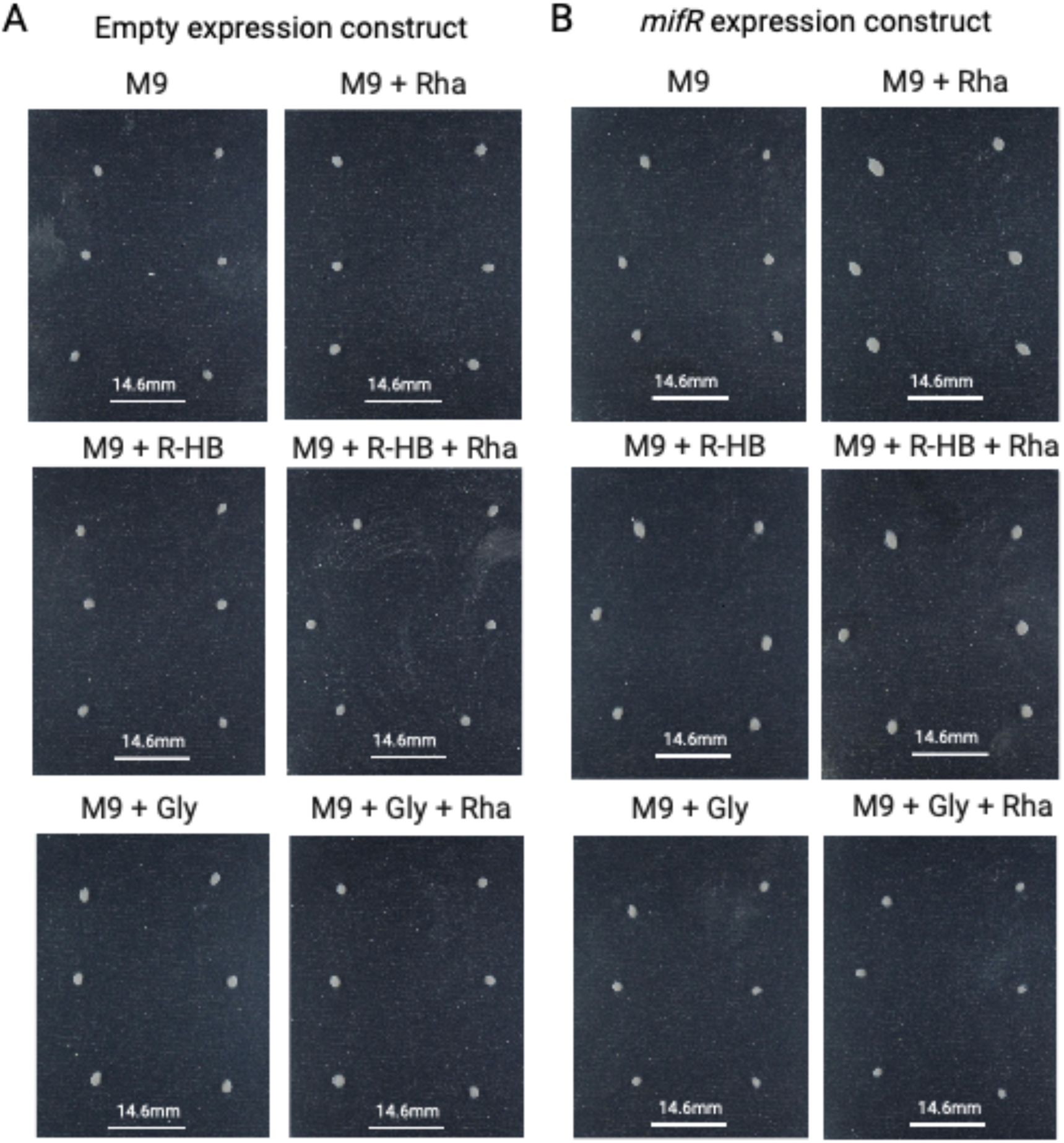
Control experiments for Fig. 4. No motility observed after 24hrs incubation for empty expression system (miniTn7[rhaSR-PrhaBAD]) or the mifR overexpression construct in M9 supplement with any combination of R-HB, Gly or Rha supplement. Confirms motility observed in Fig 3 is due to *hbcR* expression in presence of R-HB and *gcsR* expression in presence of L-glycine.

## References

1. Adhikari, S., Erill, I., Curtis, P.D., 2021. Transcriptional rewiring of the GcrA/CcrM bacterial epigenetic regulatory system in closely related bacteria. PLoS Genetics 17, 1–30. https://doi.org/10.1371/journal.pgen.1009433

2. Alhindi, T., Zhang, Z., Ruelens, P., Coenen, H., Degroote, H., Iraci, N., Geuten, K., 2017. Protein interaction evolution from promiscuity to specificity with reduced flexibility in an increasingly complex network. Scientific Reports 7, 1–15. https://doi.org/10.1038/srep44948

3. Alsohim, A.S., Taylor, T.B., Barrett, G.A., Gallie, J., Zhang, X.X., Altamirano-Junqueira, A.E., Johnson, L.J., Rainey, P.B., Jackson, R.W., 2014. The biosurfactant viscosin produced by Pseudomonas fluorescensSBW25 aids spreading motility and plant growth promotion. Environmental Microbiology 16, 2267–2281. https://doi.org/10.1111/1462-2920.12469

4. Bandyopadhyay, A., Banik, S.K., 2012. Positive feedback and temperature mediated molecular switch controls differential gene regulation in Bordetella pertussis. BioSystems 110, 107–118. https://doi.org/10.1016/j.biosystems.2012.08.004

5. Baumstark, R., Hänzelmann, S., Tsuru, S., Schaerli, Y., Francesconi, M., Mancuso, F.M., Castelo, R., Isalan, M., 2015. The propagation of perturbations in rewired bacterial gene networks. Nature Communications 6, 1–5. https://doi.org/10.1038/ncomms10105

6. Bouteiller, M., Dupont, C., Bourigault, Y., Latour, X., Barbey, C., Konto-Ghiorghi, Y., Merieau, A., 2021. Pseudomonas Flagella: Generalities and Specificities. IJMS 22, 3337. https://doi.org/10.3390/ijms22073337

7. Browning, D.F., Butala, M., Busby, S.J.W., 2019. Bacterial Transcription Factors: Regulation by Pick “N” Mix. Journal of Molecular Biology 431, 4067–4077. https://doi.org/10.1016/j.jmb.2019.04.011

8. Bush, M., Dixon, R., 2012. The Role of Bacterial Enhancer Binding Proteins as Specialized Activators of σ ^54^ -Dependent Transcription. Microbiol Mol Biol Rev 76, 497–529. https://doi.org/10.1128/MMBR.00006-12

9. Chen, S., Zhou, Y., Chen, Y., Gu, J., 2018. fastp: an ultra-fast all-in-one FASTQ preprocessor. Bioinformatics 34, i884–i890. https://doi.org/10.1093/bioinformatics/bty560

10. Choi, K.H., Schweizer, H.P., 2006. mini-Tn7 insertion in bacteria with single attTn7 sites: Example Pseudomonas aeruginosa. Nature Protocols 1, 153–161. https://doi.org/10.1038/nprot.2006.24

11. Copley, S.D., 2020. The physical basis and practical consequences of biological promiscuity. Physical Biology 17. https://doi.org/10.1088/1478-3975/ab8697

12. Copley, S.D., 2015. An evolutionary biochemist’s perspective on promiscuity. Trends in Biochemical Sciences 40, 72–78. https://doi.org/10.1016/j.tibs.2014.12.004

13. Dasgupta, N., Wolfgang, M.C., Goodman, A.L., Arora, S.K., Jyot, J., Lory, S., Ramphal, R., 2003. A four-tiered transcriptional regulatory circuit controls flagellar biogenesis in Pseudomonas aeruginosa. Molecular Microbiology 50, 809–824. https://doi.org/10.1046/j.1365-2958.2003.03740.x

14. Deatherage, D.E., Barrick, J.E., 2014. Identification of Mutations in Laboratory-Evolved Microbes from Next-Generation Sequencing Data Using breseq, in: Sun, L., Shou, W. (Eds.), Engineering and Analyzing Multicellular Systems: Methods and Protocols. Springer New York, New York, NY, pp. 165–188. https://doi.org/10.1007/978-1-4939-0554-6_12

15. diCenzo, G.C., Sharthiya, H., Nanda, A., Zamani, M., Finan, T.M., 2017. PhoU allows rapid adaptation to high phosphate concentrations by modulating PstSCAB transport rate in Sinorhizobium meliloti. Journal of Bacteriology 199, 1–20. https://doi.org/10.1128/JB.00143-17

16. Doucleff, M., Pelton, J.G., Lee, P.S., Nixon, B.T., Wemmer, D.E., 2007. Structural Basis of DNA Recognition by the Alternative Sigma-factor, σ54. Journal of Molecular Biology 369, 1070– 1078. https://doi.org/10.1016/j.jmb.2007.04.019

17. Ernst, D.C., Downs, D.M., 2016. 2-Aminoacrylate Stress Induces a Context-Dependent Glycine Requirement in *ridA* Strains of Salmonella enterica. J Bacteriol 198, 536–543. https://doi.org/10.1128/JB.00804-15

18. Fang, X., Sastry, A., Mih, N., Kim, D., Tan, J., Yurkovich, J.T., Lloyd, C.J., Gao, Y., Yang, L., Palsson, B.O., 2017. Global transcriptional regulatory network for Escherichia coli robustly connects gene expression to transcription factor activities. Proceedings of the National Academy of Sciences of the United States of America 114, 10286–10291. https://doi.org/10.1073/pnas.1702581114

19. Friedlander, T., Prizak, R., Guet, C.C., Barton, N.H., Tkačik, G., 2016. Intrinsic limits to gene regulation by global crosstalk. Nat Commun 7, 12307. https://doi.org/10.1038/ncomms12307

20. Galperin, M.Y., 2006. Structural classification of bacterial response regulators: Diversity of output domains and domain combinations. Journal of Bacteriology 188, 4169–4182. https://doi.org/10.1128/JB.01887-05

21. Goulian, M., 2010. Two-component signaling circuit structure and properties. Current Opinion in Microbiology 13, 184–189. https://doi.org/10.1016/j.mib.2010.01.009

22. Groisman, E.A., 2016. Feedback Control of Two-Component Regulatory Systems. Annual Review of Microbiology 70, 103–124. https://doi.org/10.1146/annurev-micro-102215-095331

23. Hervás, A.B., Canosa, I., Little, R., Dixon, R., Santero, E., 2009. NtrC-dependent regulatory network for nitrogen assimilation in Pseudomonas putida. Journal of Bacteriology 191, 6123–6135. https://doi.org/10.1128/JB.00744-09

24. Horton, J.S., Flanagan, L.M., Jackson, R.W., Priest, N.K., Taylor, T.B., 2021. A mutational hotspot that determines highly repeatable evolution can be built and broken by silent genetic changes. Nature Communications 12. https://doi.org/10.1038/s41467-021-26286-9

25. Igler, C., Lagator, M., Tkačik, G., Bollback, J.P., Guet, C.C., 2018. Evolutionary potential of transcription factors for gene regulatory rewiring. Nature Ecology and Evolution 2, 1633– 1643. https://doi.org/10.1038/s41559-018-0651-y

26. Isalan, M., Lemerle, C., Michalodimitrakis, K., Horn, C., Beltrao, P., Raineri, E., Garriga-Canut, M., Serrano, L., 2008. Evolvability and hierarchy in rewired bacterial gene networks. Nature 452, 840–845. https://doi.org/10.1038/nature06847

27. Joly, N., Engl, C., Jovanovic, G., Huvet, M., Toni, T., Sheng, X., Stumpf, M.P.H., Buck, M., 2010. Managing membrane stress: the phage shock protein (Psp) response, from molecular mechanisms to physiology. FEMS Microbiol Rev 34, 797–827. https://doi.org/10.1111/j.1574-6976.2010.00240.x

28. Jones, P., Binns, D., Chang, H.-Y., Fraser, M., Li, W., McAnulla, C., McWilliam, H., Maslen, J., Mitchell, A., Nuka, G., Pesseat, S., Quinn, A.F., Sangrador-Vegas, A., Scheremetjew, M., Yong, S.-Y., Lopez, R., Hunter, S., 2014. InterProScan 5: genome-scale protein function classification. Bioinformatics 30, 1236–1240. https://doi.org/10.1093/bioinformatics/btu031

29. Jothi, R., Balaji, S., Wuster, A., Grochow, J.A., Gsponer, J., Przytycka, T.M., Aravind, L., Babu, M.M., 2009. Genomic analysis reveals a tight link between transcription factor dynamics and regulatory network architecture. Molecular Systems Biology 5. https://doi.org/10.1038/msb.2009.52

30. Lamrabet, O., Plumbridge, J., Martin, M., Lenski, R.E., Schneider, D., Hindre, T., 2019. Plasticity of promoter-core sequences allows bacteria to compensate for the loss of a key global regulatory gene. Molecular Biology and Evolution 36, 1121–1133. https://doi.org/10.1093/molbev/msz042

31. Lundgren, B.R., Harris, J.R., Sarwar, Z., Scheel, R.A., Nomura, C.T., 2015. The metabolism of (R)-3-hydroxybutyrate is regulated by the enhancer-binding protein PA2005 and the alternative sigma factor RpoN in Pseudomonas aeruginosa PAO1. Microbiology 161, 2232–2242. https://doi.org/10.1099/mic.0.000163

32. Mainiero, M., Goerke, C., Geiger, T., Gonser, C., Herbert, S., Wolz, C., 2010. Differential target gene activation by the Staphylococcus aureus two-component system saeRS. Journal of Bacteriology 192, 613–623. https://doi.org/10.1128/JB.01242-09

33. Manni, M., Berkeley, M.R., Seppey, M., Simão, F.A., Zdobnov, E.M., 2021. BUSCO Update: Novel and Streamlined Workflows along with Broader and Deeper Phylogenetic Coverage for Scoring of Eukaryotic, Prokaryotic, and Viral Genomes. Molecular Biology and Evolution 38, 4647–4654. https://doi.org/10.1093/molbev/msab199

34. Martchenko, M., Levitin, A., Hogues, H., Nantel, A., Whiteway, M., 2007. Transcriptional Rewiring of Fungal Galactose-Metabolism Circuitry. Current Biology 17, 1007–1013. https://doi.org/10.1016/j.cub.2007.05.017

35. Martínez-Antonio, A., Collado-Vides, J., 2003. Identifying global regulators in transcriptional regulatory networks in bacteria. Current Opinion in Microbiology 6, 482–489. https://doi.org/10.1016/j.mib.2003.09.002

36. Meisner, J., Goldberg, J.B., 2016. The Escherichia coli rhaSR-PrhaBAD inducible promoter system allows tightly controlled gene expression over a wide range in Pseudomonas aeruginosa. Applied and Environmental Microbiology 82, 6715–6727. https://doi.org/10.1128/AEM.02041-16

37. Mistry, J., Chuguransky, S., Williams, L., Qureshi, M., Salazar, G.A., Sonnhammer, E.L.L., Tosatto, S.C.E., Paladin, L., Raj, S., Richardson, L.J., Finn, R.D., Bateman, A., 2021. Pfam: The protein families database in 2021. Nucleic Acids Research 49, D412–D419. https://doi.org/10.1093/nar/gkaa913

38. Moskowitz, S.M., Brannon, M.K., Dasgupta, N., Pier, M., Sgambati, N., Miller, A.K., Selgrade, S.E., Miller, S.I., Denton, M., Conway, S.P., Johansen, H.K., Høiby, N., 2012. PmrB mutations promote polymyxin resistance of Pseudomonas aeruginosa isolated from colistin-treated cystic fibrosis patients. Antimicrobial Agents and Chemotherapy 56, 1019–1030. https://doi.org/10.1128/AAC.05829-11

39. Nie, X., Dong, W., Yang, C., 2019. Genomic reconstruction of σ54 regulons in Clostridiales. BMC Genomics 20, 565. https://doi.org/10.1186/s12864-019-5918-4

40. Okamura-Ikeda, K., Ohmura, Y., Fujiwara, K., Motokawa, Y., 1993. Cloning and nucleotide sequence of the gcv operon encoding the Escherichia coli glycine-cleavage system. Eur J Biochem 216, 539–548. https://doi.org/10.1111/j.1432-1033.1993.tb18172.x

41. Olaitan, A.O., Morand, S., Rolain, J.-M., 2014. Mechanisms of polymyxin resistance: acquired and intrinsic resistance in bacteria. Frontiers in Microbiology 5, 643. https://doi.org/10.3389/fmicb.2014.00643

42. O’Leary, N.A., Wright, M.W., Brister, J.R., Ciufo, S., Haddad, D., McVeigh, R., Rajput, B., Robbertse, B., Smith-White, B., Ako-Adjei, D., Astashyn, A., Badretdin, A., Bao, Y., Blinkova, O., Brover, V., Chetvernin, V., Choi, J., Cox, E., Ermolaeva, O., Farrell, C.M., Goldfarb, T., Gupta, T., Haft, D., Hatcher, E., Hlavina, W., Joardar, V.S., Kodali, V.K., Li, W., Maglott, D., Masterson, P., McGarvey, K.M., Murphy, M.R., O’Neill, K., Pujar, S., Rangwala, S.H., Rausch, D., Riddick, L.D., Schoch, C., Shkeda, A., Storz, S.S., Sun, H., Thibaud-Nissen, F., Tolstoy, I., Tully, R.E., Vatsan, A.R., Wallin, C., Webb, D., Wu, W., Landrum, M.J., Kimchi, A., Tatusova, T., DiCuccio, M., Kitts, P., Murphy, T.D., Pruitt, K.D., 2016. Reference sequence (RefSeq) database at NCBI: current status, taxonomic expansion, and functional annotation. Nucleic Acids Res 44, D733–D745. https://doi.org/10.1093/nar/gkv1189

43. Patel, V., Matange, N., 2021. Adaptation and compensation in a bacterial gene regulatory network evolving under antibiotic selection. eLife 10, 1–27. https://doi.org/10.7554/eLife.70931

44. Payne, J.L., Wagner, A., 2019. The causes of evolvability and their evolution. Nat Rev Genet 20, 24–38. https://doi.org/10.1038/s41576-018-0069-z

45. Payne, J.L., Wagner, A., 2014. The Robustness and Evolvability of Transcription Factor Binding Sites. Science 343, 875–877. https://doi.org/10.1126/science.1249046

46. Pougach, K., Voet, A., Kondrashov, F.A., Voordeckers, K., Christiaens, J.F., Baying, B., Benes, V., Sakai, R., Aerts, J., Zhu, B., Van Dijck, P., Verstrepen, K.J., 2014. Duplication of a promiscuous transcription factor drives the emergence of a new regulatory network. Nature Communications 5, 1–11. https://doi.org/10.1038/ncomms5868

47. Rao, S.D., Igoshin, O.A., 2021. Overlaid positive and negative feedback loops shape dynamical properties of PhoPQ two-component system. PLoS Computational Biology 17, 1–18. https://doi.org/10.1371/journal.pcbi.1008130

48. Sarwar, Z., Lundgren, B.R., Grassa, M.T., Wang, M.X., Gribble, M., Moffat, J.F., Nomura, C.T., 2016. GcsR, a TyrR-Like Enhancer-Binding Protein, Regulates Expression of the Glycine Cleavage System in Pseudomonas aeruginosa PAO1. mSphere 1, e00020–16. https://doi.org/10.1128/mSphere.00020-16

49. Schmutzer, M., Wagner, A., 2020. Gene expression noise can promote the fixation of beneficial mutations in fluctuating environments. PLoS Comput Biol 16, e1007727. https://doi.org/10.1371/journal.pcbi.1007727

50. Schultheisz, H.L., Szymczyna, B.R., Scott, L.G., Williamson, J.R., 2011. Enzymatic De Novo Pyrimidine Nucleotide Synthesis. J. Am. Chem. Soc. 133, 297–304. https://doi.org/10.1021/ja1059685

51. Seshasayee, A.S., Bertone, P., Fraser, G.M., Luscombe, N.M., 2006. Transcriptional regulatory networks in bacteria: from input signals to output responses. Current Opinion in Microbiology 9, 511–519. https://doi.org/10.1016/j.mib.2006.08.007

52. Shepherd, M., Horton, J.S., Taylor, T.B., 2022a. A near-deterministic mutational hotspot in Pseudomonas fluorescens is constructed by multiple interacting genomic features. Molecular Biology and Evolution 39, 1–7. https://doi.org/10.1093/molbev/msac132

53. Shepherd, M., Pierce, A.P., Taylor, T.B., 2022b. Evolutionary innovation through transcription factor promiscuity in microbes is constrained by pre-existing gene regulatory network architecture. https://doi.org/10.1101/2022.07.12.499626

54. Silby, M.W., Cerdeño-Tárraga, A.M., Vernikos, G.S., Giddens, S.R., Jackson, R.W., Preston, G.M., Zhang, X.X., Moon, C.D., Gehrig, S.M., Godfrey, S.A.C., Knight, C.G., Malone, J.G., Robinson, Z., Spiers, A.J., Harris, S., Challis, G.L., Yaxley, A.M., Harris, D., Seeger, K., Murphy, L., Rutter, S., Squares, R., Quail, M.A., Saunders, E., Mavromatis, K., Brettin, T.S., Bentley, S.D., Hothersall, J., Stephens, E., Thomas, C.M., Parkhill, J., Levy, S.B., Rainey, P.B., Thomson, N.R., 2009. Genomic and genetic analyses of diversity and plant interactions of Pseudomonas fluorescens. Genome Biology 10, 1–16. https://doi.org/10.1186/gb-2009-10-5-r51

55. Studholme, D.J., Buck, M., 2000. The biology of enhancer-dependent transcriptional regulation in bacteria: insights from genome sequences. FEMS Microbiology Letters 186, 1–9. https://doi.org/10.1111/j.1574-6968.2000.tb09074.x

56. Taylor, T., Mulley, G., McGuffin, L., Johnson, L., Brockhurst, M., Arseneault, T., Silby, M., Jackson, R., 2015b. Evolutionary rewiring of bacterial regulatory networks. Microbial Cell 2, 256–258. https://doi.org/10.15698/mic2015.07.215

57. Taylor, T.B., Mulley, G., Dills, A.H., Alsohim, A.S., McGuffin, L.J., Studholme, D.J., Silby, M.W., Brockhurst, M.A., Johnson, L.J., Jackson, R.W., 2015a. Evolutionary resurrection of flagellar motility via rewiring of the nitrogen regulation system. Science 347, 1014–1017. https://doi.org/10.1126/science.1259145

58. Taylor, T.B., Shepherd, M., Jackson, R.W., Silby, M.W., 2022. Natural selection on crosstalk between gene regulatory networks facilitates bacterial adaptation to novel environments. Current Opinion in Microbiology 67, 102140. https://doi.org/10.1016/j.mib.2022.02.002

59. Tirosh, I., Barkai, N., Verstrepen, K.J., 2009. Promoter architecture and the evolvability of gene expression. J Biol 8, 95. https://doi.org/10.1186/jbiol204

60. Tsuda, M.E., Kawata, M., 2010. Evolution of gene regulatory networks by fluctuating selection and intrinsic constraints. PLoS Computational Biology 6. https://doi.org/10.1371/journal.pcbi.1000873

61. West, T.P., 2010. Effect of carbon source on pyrimidine biosynthesis in *Pseudomonas oryzihabitans*: Effect of carbon source on pyrimidine biosynthesis in *Pseudomonas oryzihabitans*. J. Basic Microbiol. 50, 397–400. https://doi.org/10.1002/jobm.201000022

62. Weyder, M., Prudhomme, M., Bergé, M., Polard, P., Fichant, G., 2018. Dynamic modeling of Streptococcus pneumoniae competence provides regulatory mechanistic insights into its tight temporal regulation. Frontiers in Microbiology 9, 1–25. https://doi.org/10.3389/fmicb.2018.01637

63. Yan, Q., Rogan, C.J., Pang, Y.-Y., Davis, E.W., Anderson, J.C., 2020. Ancient co-option of an amino acid ABC transporter locus in Pseudomonas syringae for host signal-dependent virulence gene regulation. PLoS Pathog 16, e1008680. https://doi.org/10.1371/journal.ppat.1008680

64. Zhou, T., Huang, J., Liu, Z., Xu, Z., Zhang, L., 2021. Molecular Mechanisms Underlying the Regulation of Biofilm Formation and Swimming Motility by FleS/FleR in Pseudomonas aeruginosa. Front. Microbiol. 12, 707711. https://doi.org/10.3389/fmicb.2021.707711

